# GlycoEnzDB: A database of enzymes involved in human glycosylation

**DOI:** 10.1101/2025.08.30.673257

**Authors:** Yusen Zhou, Vishnu Ghosh, Shriramprasad Venkatesan, Shyam Sriram, Edward Sobczak, Srirangaraj Setlur, Rudiyanto Gunawan, Sriram Neelamegham

## Abstract

The glycan distribution on cells is governed by the stochastic activity of different families of enzymes that are together called ‘glycoEnzymes’. These include ~400 gene products or 2% of the proteome, that have recently been curated in an ontology called GlycoEnzOnto. With the goal of making this ontology more accessible to the larger biomedical and biotechnology community, we organized a web resource called GlycoEnzDB, presenting this enzyme classification both in terms of enzyme function and the pathways that they participate in. This information is linked to i) Figures from the “Essentials of Glycobiology” textbook, ii) General gene, enzyme and pathway data appearing in external databases, iii) Manual and generative-artificial intelligence (AI) based text describing the function and pathways regulated by these entities, iv) Single-cell expression data across cell lines, normal human cell-types and tissue, and v) CRISPR-knockout/activation/inactivation and Transcription factor activity predictions. Whereas these data are curated for human glycoEnzymes, the knowledge framework may be extended to other species also. The user–friendly web interface is accessible at www.virtualglycome.org/glycoenzdb.

## INTRODUCTION

Glycosylation is a ubiquitous post-translational modification in mammals that commonly results in the formation of complex carbohydrates or glycans, on diverse protein and lipid scaffolds, mediating numerous biological functions (Varki, A. 2017). The formation of glycan structures is driven by the action of ~400 genes that are called ‘glycoGenes’ (Neelamegham, S. and Mahal, L.K. 2016). Each of these glycoGenes is considered to produce corresponding ‘glycoEnzymes’, even though not all molecules strictly display enzymatic activity. These glycoEnzymes, which constitute ~2% of the expressed proteome, have recently been classified based on an ontology called ‘GlycoEnzOnto’ (Groth, T., Diehl, A.D., et al. 2022). Such semantic literature-based organization of knowledge allows for the description of interactions among these molecules. The curation in GlycoEnzOnto is based on the Gene Ontology convention (Ashburner, M., Ball, C.A., et al. 2000), where each entry is described in terms of its: i) function or molecular role, ii) pathway or biological process it regulates, and iii) localization or physical location in or outside cells. As glycoEnzymes largely lie in the endoplasmic reticulum or Golgi with a few involved in lysosomal degradation, much of the focus of this manuscript is on ‘function’ and ‘pathway’ relations.

The goal of this manuscript is to introduce a resource called GlycoEnzDB that is freely available as part of the virtualglycome.org resource. The effort is distinct from previous work in the field. Notably, the GlycoGene Database (GGDB), which is now part of the GlyCosmos portal (Yamada, I., Shiota, M., et al. 2020), curates a limited set of glycosylation genes. The gene set encompassed by GlycoEnzDB is more comprehensive and it is presented as an extension of a well-defined ontology. The Moremen laboratory has curated glycosylation pathways in mammals but these pathways are only associated with limited functional data, primarily focused on recombinant DNA technology and enzyme production (https://glycoenzymes.ccrc.uga.edu/, (Moremen, K.W., Tiemeyer, M., et al. 2012)). Glycopacity (https://glyco.me/) uses gene expression to assess the capacity of individual glycosylation pathways to produce glycans along specified pathways (Dworkin, L.A., Clausen, H., et al. 2022, Schjoldager, K.T., Narimatsu, Y., et al. 2020). In contrast, this paper does not focus on capacity, but rather attempts to link glycobiological knowledge with textbooks in the field, expression data in cell lines and single-cells, and also estimates related to CRISPR sgRNA efficiency and putative transcription factor-glycogene relations.

The focus of GlycoEnzDB is both on new investigators in the field attempting to grasp glycosylation pathways and more experienced scientists looking for well-organized experimental data. One of its goals is to provide an intuitive interface that would be helpful to experimentalists in the biochemistry and biomedical sciences fields. Another goal is to provide a platform for collating enzyme-centric data related to glycosylation pathways. As a starting point, a limited set of experimental data are curated in the current GlycoEnzDB framework, and we expect to expand on this in the future based on our ongoing research in our and other laboratories.

## RESULTS AND DISCUSSION

The glycoEnzymes are an important starting point for studies of glycobiology, as they define the reaction pathways used to produce glycans. GlycoEnzDB curates 403 human glycoEnzymes based on their function (**Supplemental Table S1**) and the pathways (**Supplemental Table S2**) they participate in. From the “function” perspective, the glycoEnzymes are grouped into glycosyltransferases (sialyltransferases, fucosyltransferases, etc.), other transferases (e.g., sulfotransferases), modifying enzymes (e.g., epimerases and kinases), glycosidases, molecular transporters, and other regulators. Additionally, the glycoEnzymes are also classified according to their “pathways” in terms of their roles in glycoconjugate biosynthesis, degradation, donor synthesis, and transport. These biosynthesis pathways include the steps involved in: (i) ‘Core’ biosynthesis that results in the attachment of the first monosaccharide(s) or oligosaccharide to the protein/lipid that initiates the biosynthesis of specific glycoconjugate types. In this regard, the core pathways initiate the biosynthesis of glycolipids (glycosphingolipid and GPI pathways), glycosaminoglycans (hyaluronic acid, chondroitin/dermatan sulfate, heparan sulfate), O-linked glycans (O-GalNAc type, O-GlcNAc and additional rarer modifications), C-Mannosyl linkages at Trp, and N-linked glycans commonly at the Asn-X-Ser/Thr sequon; (ii) ‘Elongation and branching’ reactions that extend the original glycan via the extension of chondroitin/dermatan, heparan, and keratan sulfate chains, LacNAc (N-acetyl lactosamine), LacdiNAc (GalNAc(β1-4)GlcNAcβ) and I-branching (Gal(1-4)[GlcNAc(β1-6)]Galβ); and (iii) ‘Termination or capping’ processes that often involve sulfation, fucosylation and sialylation reactions.

Many of the above functions and pathways are linked to 24 figures that are adapted from the Essentials of Glycobiology (EoG) textbook that capture the human glycosylation biosynthesis process (Varki, A., Cummings, R.D., et al. 2022). **Figure 1A** presents a screenshot of the front page of the database. Clicking on the “Function” and “Pathways” buttons displays a list of genes involved in the function/pathway based on GlycoEnzOnto. A relevant figure from the EoG text is presented in many cases. The enzyme names in these figures appear in red, and clicking on any name will take the user to more detailed information pages described below. Additionally, if enzymes are searched based on their HGNC gene name, this also opens the details page for that enzyme along with corresponding EoG figure(s), when available. To better explain the enzyme activity, we have manually curated text related to individual enzyme function. This is primarily collated from the EoG textbook (Varki, A., Cummings, R.D., et al. 2022) and the “Handbook of glycosyltransferases and related genes” (Taniguchi, N., Honke, K., et al. 2014). The EoG text was also used to train a custom GPT (Generative pre-trained transformer model) called “Dr. Glyco-GPT”. This new GPT can be queried with glycosylation related questions. For GlycoEnzDB, the GPT was asked to write a short essay about each of the curated enzymes, with respect to their: i) function, ii) reaction pathways, ii) location, and iv) disease relation. The output is presented in the database, with only limited manual curation. Dr. Glyco-GPT can also be used to answer additional user queries, but one must caution that the use of large-language models (LLMs) can be error-prone and assertions need to be validated manually against published literature. Additionally, DrawGlycan-SNFG is used to draw reaction schemes describing the catalytic activity of the individual glycoEnzymes (Cheng, K., Pawlowski, G., et al. 2019, Cheng, K., Zhou, Y., et al. 2017). This uses the Symbol Nomenclature for Glycans (SNFG) nomenclature for rendering pictorial schematics of complex carbohydrates (Neelamegham, S., Aoki-Kinoshita, K., et al. 2019, Varki, A., Cummings, R.D., et al. 2015).

**Figure 1.**
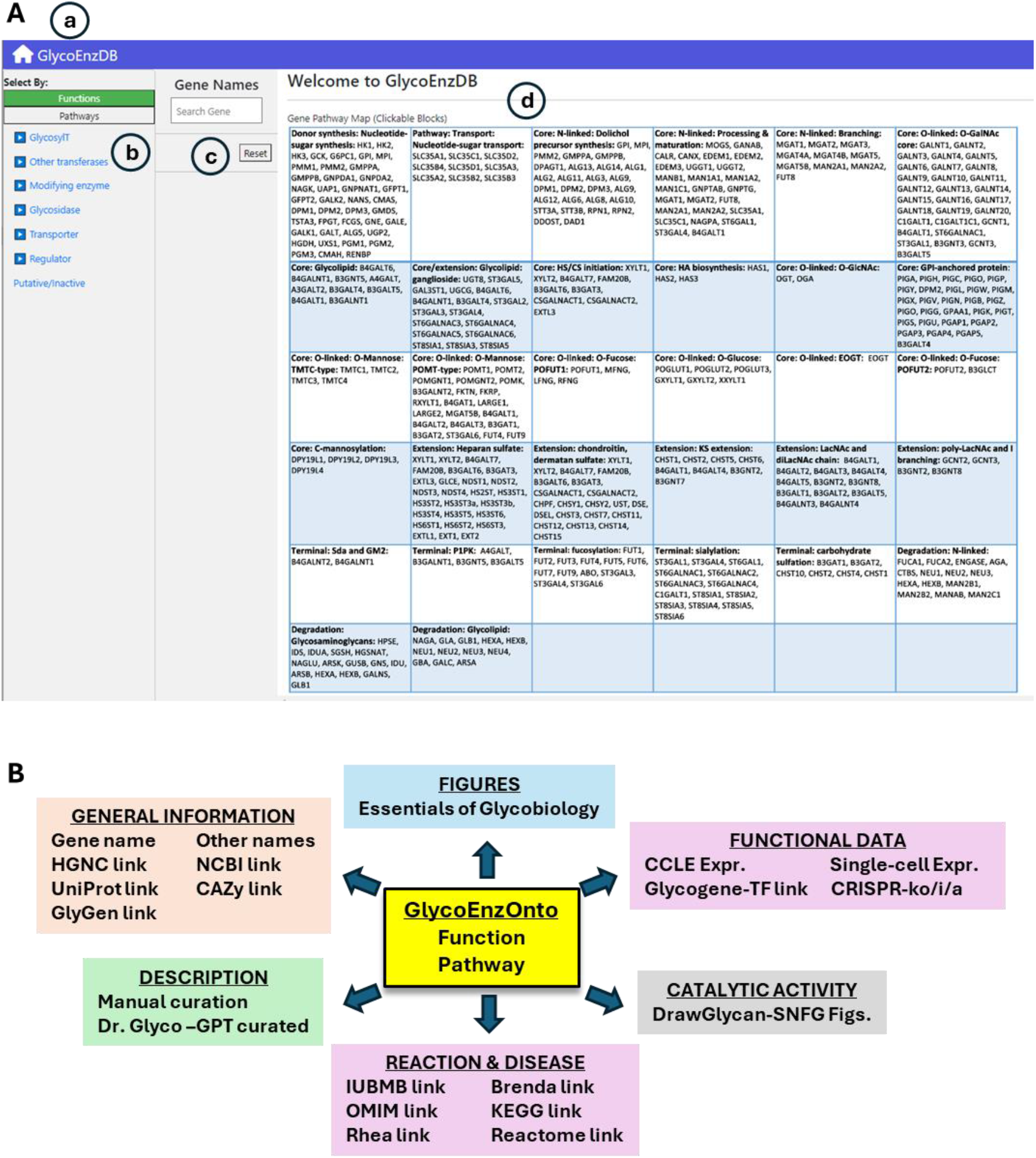
GlycoEnzDB: **A**. Landing page for web resource. Icons highlight: **a)** Home link to return to this page; **b)** Pull down menus to parse GlycoGenes/Enzymes based on either function or pathway; **c)** Search field based on gene HGNC name; **d)** 32 mammalian glycosylation pathways curated from the Essentials of Glycobiology textbook. **B**. Other resources and links available at GlycoEnzDB database.

Besides manual curation, GlycoEnzDB also integrates information for each enzyme using external databases listed in **Table 1**. These include resources that provide community-consensus HGNC gene names (Seal, R.L., Braschi, B., et al. 2023), and databases related to nucleic acids (NCBI: (Haft, D.H., Badretdin, A., et al. 2024)), proteins (UniProt: (UniProt, C. 2023)), glycans (GlyGen: (York, W.S., Mazumder, R., et al. 2020)), diseases (OMIM: (Amberger, J.S. and Hamosh, A. 2017)), enzymes (CAZy: (Drula, E., Garron, M.L., et al. 2022), IUBMB: (McDonald, A.G., Boyce, S., et al. 2009), Brenda: (Chang, A., Jeske, L., et al. 2021)), reaction rules (Rhea: (Bansal, P., Morgat, A., et al. 2022)) and pathway maps (KEGG: (Hashimoto, K., Goto, S., et al. 2006), Reactome: (Milacic, M., Beavers, D., et al. 2024)) (**Figure 1B**). Additional wet-lab data and computational predictions are provided to connect textbook knowledge with experiments. Currently, this is done by scraping glycogene expression data for various cancer cell lines from DepMap (Ghandi, M., Huang, F.W., et al. 2019), and single-cell glycogene data from Tabula Sapiens (Tabula Sapiens, C., Jones, R.C., et al. 2022). In this regard, DepMap includes bulk sequencing data for ~2000 cell lines including many of the cells that are part of the NCI-60 collection. The scRNA-seq (single-cell RNA-seq) data from Tabula Sapiens comprises measurements from 483,152 human cells from 15 donors that are organized into 24 tissues and over 400 cell types. We also provide CRISPR knockout (CRISPR-ko), activation (CRISPR-a) and inactivation (CRISPR-i) sequences from the CRISPick resource (Doench, J.G., Fusi, N., et al. 2016). Finally, the database provides computational predictions of TFs regulating glycosylation by examining mutual information of scRNA-seq from the Tabula Sapiens dataset for every TF-glycoGene pair from the TF-links database (Chrysinas, P., Venkatesan, S., et al. 2024). **Figure 2** provides webpage vignettes for a representative enzyme.

**Table 1:**
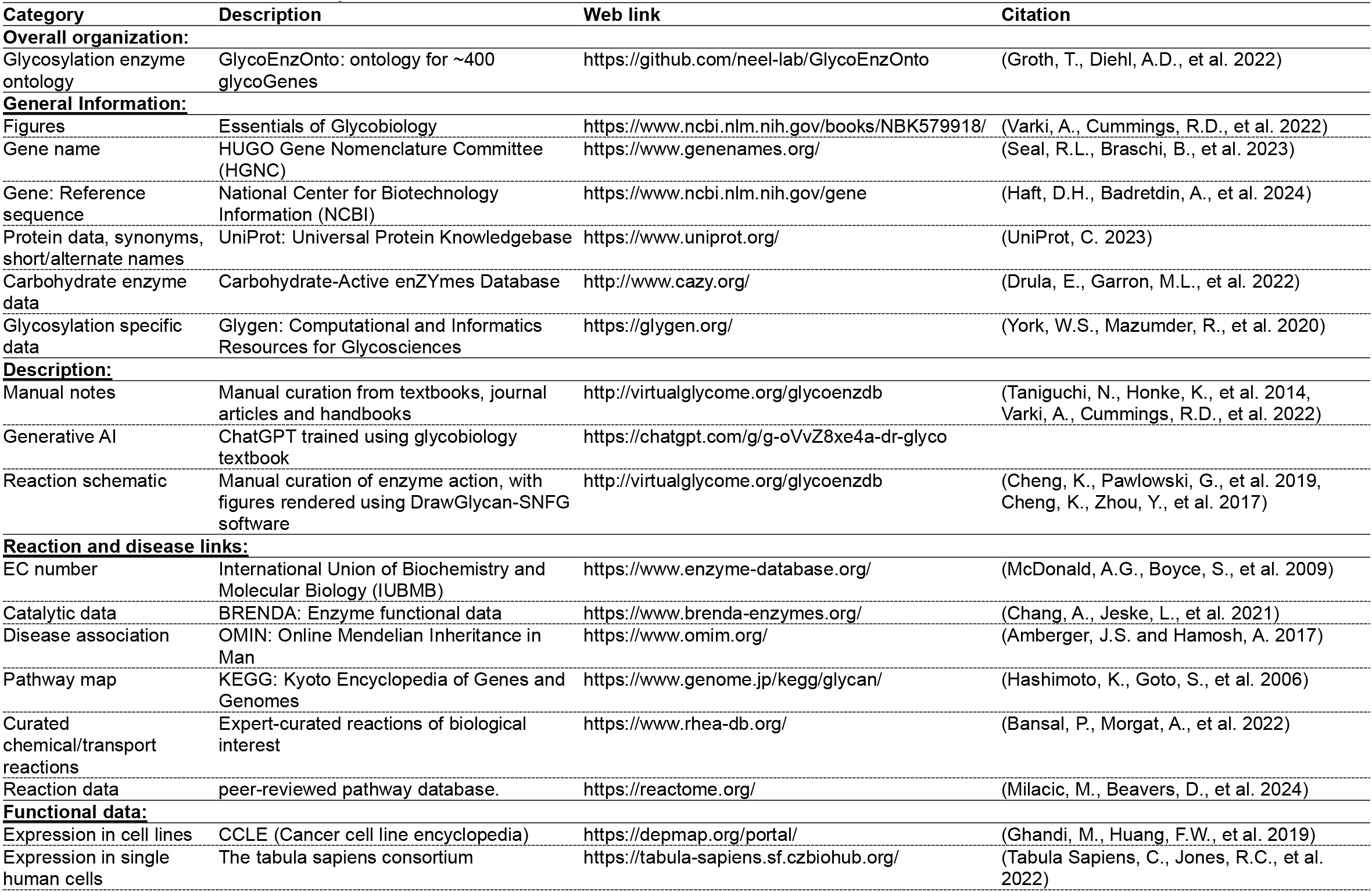
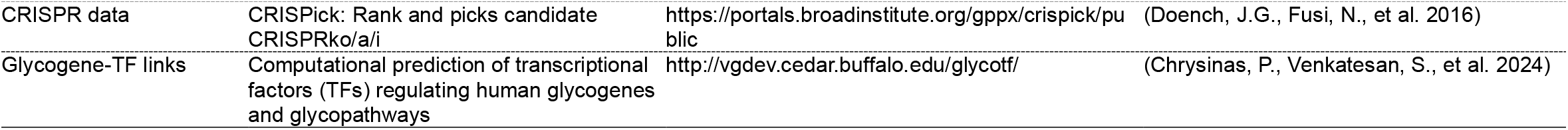
Sources of data curated at GlycoEnzDB.

| Category | Description | Web link | Citation |
| --- | --- | --- | --- |
| **Overall organization:** |  |  |  |
| Glycosylation enzyme ontology | GlycoEnzOnto: ontology for ~400 glycoGenes | https://github.com/neel-lab/GlycoEnzOnto | (Groth, T., Diehl, A.D., et al. 2022) |
| **General Information:** |  |  |  |
| Figures | Essentials of Glycobiology | https://www.ncbi.nlm.nih.gov/books/NBK579918/ | (Varki, A., Cummings, R.D., et al. 2022) |
| Gene name | HUGO Gene Nomenclature Committee (HGNC) | https://www.genenames.org/ | (Seal, R.L., Braschi, B., et al. 2023) |
| Gene: Reference sequence | National Center for Biotechnology Information (NCBI) | https://www.ncbi.nlm.nih.gov/gene | (Haft, D.H., Badretdin, A., et al. 2024) |
| Protein data, synonyms, short/alternate names | UniProt: Universal Protein Knowledgebase | https://www.uniprot.org/ | (UniProt, C. 2023) |
| Carbohydrate enzyme data | Carbohydrate-Active enZymes Database | http://www.cazy.org/ | (Drula, E., Garron, M.L., et al. 2022) |
| Glycosylation specific data | Glygen: Computational and Informatics Resources for Glycosciences | https://glygen.org/ | (York, W.S., Mazumder, R., et al. 2020) |
| **Description:** |  |  |  |
| Manual notes | Manual curation from textbooks, journal articles and handbooks | http://virtualglycome.org/glycoenzdb | (Taniguchi, N., Honke, K., et al. 2014, Varki, A., Cummings, R.D., et al. 2022) |
| Generative AI | ChatGPT trained using glycobiology textbook | https://chatgpt.com/g/g-oVvZ8xe4a-dr-glyco |  |
| Reaction schematic | Manual curation of enzyme action, with figures rendered using DrawGlycan-SNFG software | http://virtualglycome.org/glycoenzdb | (Cheng, K., Pawlowski, G., et al. 2019, Cheng, K., Zhou, Y., et al. 2017) |
| **Reaction and disease links:** |  |  |  |
| EC number | International Union of Biochemistry and Molecular Biology (IUBMB) | https://www.enzyme-database.org/ | (McDonald, A.G., Boyce, S., et al. 2009) |
| Catalytic data | BRENDA: Enzyme functional data | https://www.brenda-enzymes.org/ | (Chang, A., Jeske, L., et al. 2021) |
| Disease association | OMIM: Online Mendelian Inheritance in Man | https://www.omim.org/ | (Amberger, J.S. and Hamosh, A. 2017) |
| Pathway map | KEGG: Kyoto Encyclopedia of Genes and Genomes | https://www.genome.jp/kegg/glycan/ | (Hashimoto, K., Goto, S., et al. 2006) |
| Curated chemical/transport reactions | Expert-curated reactions of biological interest | https://www.rhea-db.org/ | (Bansal, P., Morgat, A., et al. 2022) |
| Reaction data | peer-reviewed pathway database. | https://reactome.org/ | (Milacic, M., Beavers, D., et al. 2024) |
| **Functional data:** |  |  |  |
| Expression in cell lines | CCLE (Cancer cell line encyclopedia) | https://depmap.org/portal/ | (Ghandi, M., Huang, F.W., et al. 2019) |
| Expression in single human cells | The tabula sapiens consortium | https://tabula-sapiens.sf.czbiohub.org/ | (Tabula Sapiens, C., Jones, R.C., et al. 2022) |
| CRISPR data | CRISPick: Rank and picks candidate<br>CRISPRko/a/i | https://portals.broadinstitute.org/gppx/crispick/public | (Doench, J.G., Fusi, N., et al. 2016) |
| Glycogene-TF links | Computational prediction of transcriptional<br>factors (TFs) regulating human glycogenes<br>and glycopathways | http://vgdev.cedar.buffalo.edu/glycotf/ | (Chrysinas, P., Venkatesan, S., et al. 2024) |

**Figure 2.**
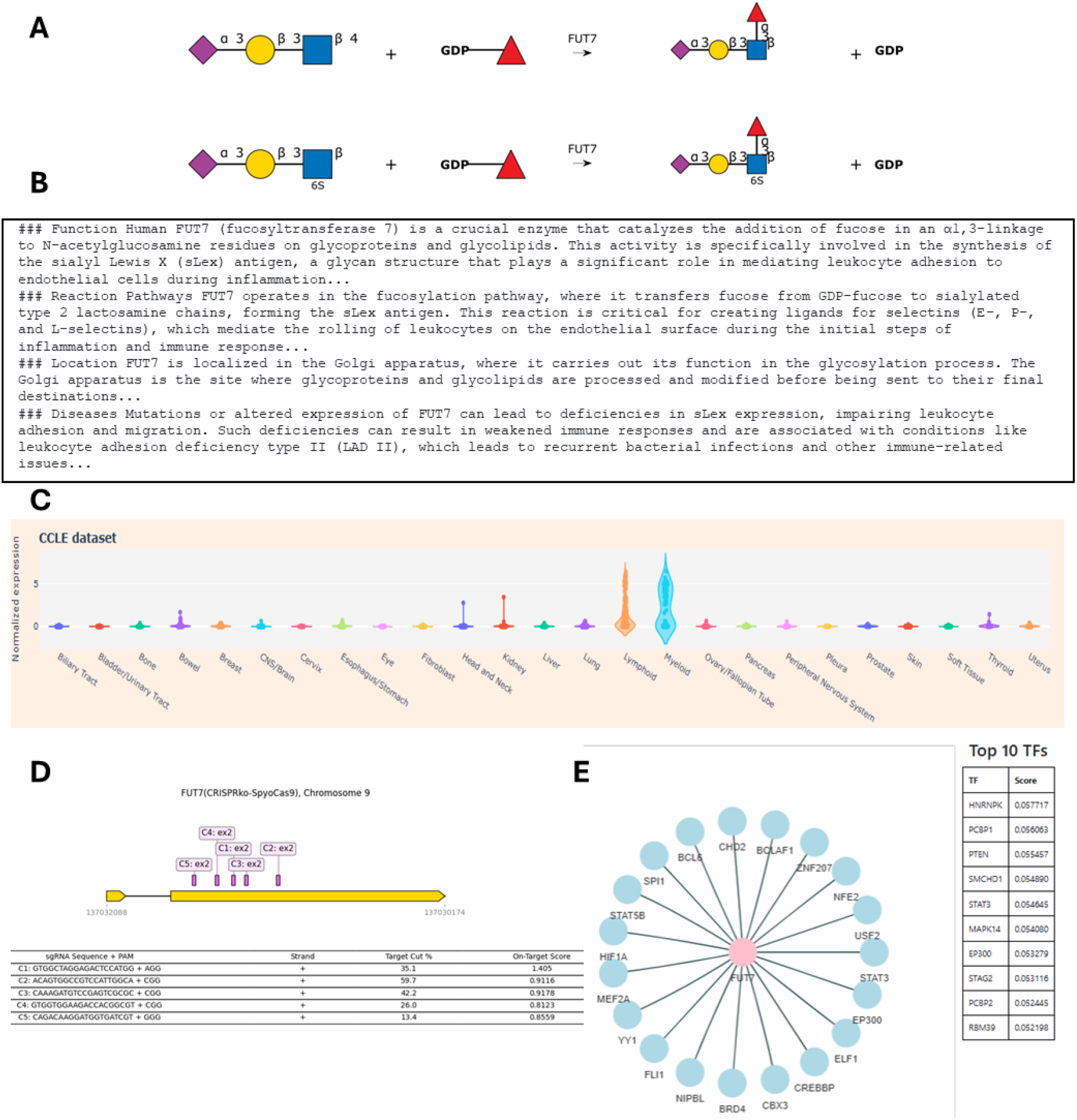
Figure vignettes: Sample data for FUT7 showing: **A**. Catalytic reaction. **B**. Sample Dr.Glyco-GPT text. **C**. Cell line transcript expression data; **D**. CRISPR knockout figure; **E**. Putative link between glycogenes and transcription factors.

Although GlycoEnzDB is curated for human glycoenzymes, many of the same or similar pathways occur in other mammals, with a few exceptions. Among these is Cytidine monophospho-N-acetylneuraminic acid hydroxylase (CMAH), which catalyzes the conversion of Neu5Ac (N-acetylneuraminic acid) to Neu5Gc (N-glycolylneuraminic acid). CMAH is present in Old World primates and many other deuterostome species but is inactive in Homo sapiens and has been independently lost in several other taxa, including the common ancestor of all birds. GGTA1 (α3Gal-T), which transfers Gal from UDP-Gal to the nonreducing terminal β-Gal of glycoconjugates to form alpha-Gal (Galα1-3Gal), is found in most non-primate mammals but is absent in Old World primates, including humans, due to gene inactivation. Finally, GBGT1, which provides α(1-3)GalNAc activity necessary to generate the Forssman glycolipid antigen, is inactive in most humans, although functional alleles persist in a minority of individuals. Scripts are available on the GlycoEnzDB GitHub repository to extend the database to non-human species.

The current version of GlycoEnzDB is focused on a limited set of experimental data. We anticipate that this will grow in the coming years with large-scale characterization of microRNA, epigenetic factors, and regulatory elements controlling glycosylation. We will endeavor to add these insights as they emerge. The enzyme rules and pathway maps provided by the resource may also facilitate the automated development of custom reaction pathway maps for model building (Huang, Y.F., Aoki, K., et al. 2021, Liu, G. and Neelamegham, S. 2015, Neelamegham, S. and Liu, G. 2011).

## MATERIALS AND METHODS

### Sources of data

The GlycoEnzDB is built on the Django python framework using the PostgreSQL relational database management system. This is an open-source resource. Source data are provided at https://github.com/neel-lab/webGlycoEnzDB, and they come from original resources listed in **Table 1**. This GitHub repository also contains code used to generate the website, data tables and accompanying figures.

### General information and enzyme description

General information about each enzyme including the gene name and links to external databases related to enzymes and pathways were gleaned from the UniProt server (UniProt, C. 2023). Notes related to enzymes were mostly manually curated from textbooks in the field. Reaction figures were rendered using the DrawGlycan-SNFG software (Cheng, K., Pawlowski, G., et al. 2019, Cheng, K., Zhou, Y., et al. 2017), based on recommended SNFG standards (Neelamegham, S., Aoki-Kinoshita, K., et al. 2019, Varki, A., Cummings, R.D., et al. 2015).

### Dr.Glyco-GPT

A custom GPT was generated using tools at ChatGPT.com. Knowledge from the EoG (Varki, A., Cummings, R.D., et al. 2022) was input used for training, with permission from the textbook editors. The generative resource was then prompted to write a 100-350 word essay about each of the enzymes with respect to their: i) function, ii) reaction pathways, ii) location, and iv) diseases. Note that, while language models such as ChatGPT are powerful tools for information synthesis, the output can be incorrect or outdated. Users, thus, need to consider the output in context of peer-reviewed literature.

### Cancer cell expression data (CCLE)

TPM values for all DepMap cell types was downloaded in April 2023 using the ‘OmicsExpressionProteinCodingGenesTPMLogp1.csv’ file. Non-glycogenes were removed and all DepMap cells were preserved. These data are plotted as violin plots. Values are inferred from RNA-seq data using the RSEM tool and are reported after log2 transformation, i.e. log2(TPM+1).

### Single-cell expression data

10X genomics based scRNA-seq data (456,101 cells) were from the Tabula Sapiens (TS) project (Tabula Sapiens, C., Jones, R.C., et al. 2022). Data were processed as described previously (Chrysinas, P., Venkatesan, S., et al. 2024). Briefly, the UMI counts for each cell were normalized to 10,000 and log transformed (log1p), to obtain relative RNA abundance data. These are plotted for the enzymes in the GlycoEnzOnto (Groth, T., Diehl, A.D., et al. 2022). Plots are generated to quantify transcript abundance in individual tissue types and also cell types or “compartments” in that tissue.

### CRISPR data

CRISPR knockout, activation and inactivation data were curated from the CRISPick resource using the human GRCh38 reference genome, and *S. pyrogenes* containing the NGG PAM sequence. Five sgRNA were chosen for each glycogene based on a previously published algorithm (Doench, J.G., Fusi, N., et al. 2016). Figures presented at the GlycoEnzDB resource were generated using the DNA_features_viewer package. They present sgRNA target location and “on-target scores” which quantify the theoretical efficacy of each of the sgRNA.

### Glycogene-transcription factor relation

TF-glycogene pairs are curated from the TFLink database, and this was previously compiled from numerous databases that relied on different evidence(s) of TF binding to regulatory elements on genes (Liska, O., Bohar, B., et al. 2022). Mutual information (MI) was used to provide additional evidence for TF-glycogene interactions based on single-cell expression data from the Tabula Sapiens. Here, MI is calculated according to:

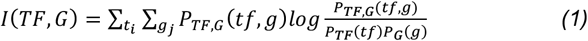

where *P*_*TF,G*_ denotes the joint probability distribution of single-cell expression of a transcription factor *TF* and a glycogene *G*, and *P*_*TF*_ and *P*_*G*_ denote the marginal probability distribution of *TF* and *G*, respectively. We evaluated *I*(*TF, G*) using the scaled UMI counts from all 10X cells in the TS dataset. More details are available elsewhere (Chrysinas, P., Venkatesan, S., et al. 2024).

## ACKNOWLEDGEMENTS

We are grateful to Theodore Groth for leading the curation of the GlycoEnzOnto ontology, and David Huang for writing scripts that parse Uniprot. We are indebted to Profs. Ajit Varki and Pamela Stanley, past and current editor-in-chief of the EoG, for supporting this work including providing comments and edits. The work would not be possible without the figures from the EoG rendered by Prof. Rick Cummings. We also thank the EoG editors and the Cold Spring Harbor Laboratory Press for allowing us to adapt the text and figures for use at GlycoEnzDB.

## DATA AVAILABILITY STATEMENT

All data, instructions and source code used to generate the web interface are available at: https://github.com/neel-lab/webGlycoEnzDB. GlycoEnzDB is deployed at: www.virtualglycome.org/glycoenzdb.

## AUTHOR CONTRIBUTIONS

Y.Z., V.G., E.S., S.S., S.V.: Methodology, Investigation; S.S.: Resources, supervision; R.G: Methodology, investigation, supervision; S.N.: Conceptualization, supervision, methodology, investigation, writing-original draft; All authors: Writing – review & editing.

